# Genomic Variation, Population Structure, and Gene Flow across Asian Pikas

**DOI:** 10.1101/2022.10.22.513340

**Authors:** Nishma Dahal, Melia G Romine, Sunita Khatiwada, Uma Ramakrishnan, Sangeet Lamichhaney

**Affiliations:** Biotechnology Division, CSIR-Institute of Himalayan Bioresource Technology, Palampur, HP, 176061, India; School of Biomedical Sciences, Kent State University, Kent, OH 44240, USA; Forest and Environment Department, Government of Sikkim, Gangtok, India; National Centre for Biological Sciences, TIFR, Bellary Road, Bangalore 560065, India; Department of Biological Sciences, Kent State University, Kent 44242, USA

## Abstract

Asian pikas have one of the most complex systematics and evolutionary history. The Himalayas is an important habitat for Asian pikas as it hosts 23 – 25% of pika’s global diversity and has provided the ancestral training ground for local adaptation to high altitudes. They are one of the most abundant species in the Himalayas and Qinghai Tibetan Plateau (QTP), however genetic studies to explore their population structure and evolution are limited. Here, we utilize a population genomics approach using ~28,000 genome-wide SNP markers obtained from restriction-site associated DNA sequencing in six species of Asian Pikas *(Ochotona* spp.) that are distributed across the Himalayas and neighboring high-altitude mountains. We examined intra- and inter-species genetic diversity, population structure, phylogenetic history and explored processes that shaped the current genetic diversity of Pikas across the Himalayas. We identified low nucleotide diversity and high inbreeding coefficient across all species which possibly indicated decreasing population size in these species. We also identified extensive evidence of gene flow (both historic and contemporary) across these species. Our findings indicate that inter-species gene flow is a key evolutionary process that has been countering the negative effect of low genetic diversity among Asian pikas.

## Introduction

The complexity of speciation was identified early on by Darwin and was referred to as the “mystery of mysteries”. Though reproductive isolation is a crucial feature of speciation, interspecies hybridization is commonly reported (Schwarz et al. 2005; Mavárez et al. 2006; Larsen et al. 2010; Hermansen et al. 2011; Kang et al. 2013; Yakimowski and Rieseberg 2014; Lamichhaney et al. 2018; Adavoudi and Pilot 2022; Combe et al. 2022). The availability of genomic data from non-model organisms has expanded our knowledge of the processes of speciation in the wild population (Storz and Hoekstra 2007; Nei and Nozawa 2011; Ravinet et al. 2017). For species living in high-altitude rugged terrain with cyclical glacial history, the process of speciation is an entangled phenomenon of isolation, reinforcement, and gene flow. The topographic alteration which results in the uplift of mountains can cause fragmentation of a continuous population, thereby causing secession of gene flow and initiating the process of speciation (Mayr 1963). Apart from topography, the climatic barrier may also aid in the divergence of the population (Dahal et al. 2017) ultimately leading to reproductive isolation. But the cyclical nature of climate may also assist in gene flow between once-isolated populations (Camurugi et al. 2021). Diverse ecological opportunities in mountains also support speciation resulting from reinforcement led by factors like selective sweeps, mutation rate variation, and recombination (Werhahn et al. 2020).

The Himalayas and the surrounding mountains, especially the eastern Himalayas, are perhaps the most rugged landscape in the world (Enkelmann et al. 2011). The Himalayas hold enormous biodiversity and these high-altitude regions are home to many unique biotas (Sharma et al. 2008). Unlike the mountains of North America and Europe, the process of glaciation and their timing appears to be much more complex in the Himalayas (Shi et al. 1992; Dimri et al. 2020), thus providing foundational opportunities to study the complexities 3 in speciation aided by topography and climatic history. The pattern of glaciation in the Himalayas was more localized (Owen et al. 2005), probably impacting the high-altitude species (Dahal et al. 2021). Further, it has also been hypothesized that glaciation in the east and the west Himalayas were asynchronous (Owen et al. 2005) resulting in current nested species diversity (Srinivasan et al. 2014). Thus, genetic studies on the high-altitude biotas of the Himalayas and surrounding mountains could potentially show interesting patterns driven by reinforcement and gene flow. Only few genetic studies (Werhahn et al. 2020; Arekar et al. 2022) have carried out comparative studies across the latitudinal gradient in the Himalayas, however, these studies have not explored the genetic diversity between east and west Himalayan populations.

Pikas are one of such high altitude, and climate change sensitive species (Smith 2020). Despite being one of the most abundant species in the high-altitude regions of Himalaya and Qinghai Tibetan Plateau (QTP), studies on these Asian pikas are limited, which may be critically important, first of all, to understand their population dynamics and also for their management and conservation (Palsboll et al. 2007). Of the six Asian pikas species reported from the Himalayas (Dahal et al. 2020), ecology and genetics of only two species – *O. curzoniae* and *O. roylei* have been explored (Bhattacharyya et al. 2013, 2014, 2019; Qu et al. 2017). However, these studies were limited to understanding the relative abundance of these species as a correlate of habitat type and climatic variables. So, with no information on population dynamics, inferences from population genetic studies on Asian pikas can provide first-hand information on the underlying impacts of ongoing climate change, which is reported to be extreme in this part of their range (Shrestha et al. 2012).

The Himalayas and the Tibetan plateau form major distribution centers for at least two species-rich subgenera (*Conothoa* and *Ochotona*) out of a total of five recognized subgenera (Wang et al. 2020; Lissovsky et al. 2022) of Pikas. Few studies (Dahal et al. 2017; Lissovsky et al. 2017, 2019, 2022; Koju et al. 2017) have sampled pikas across the Himalayas, mostly to resolve the taxonomic ambiguities. Two studies have described the distinct lineages of pikas, both sampled from eastern Himalayas (Dahal et al. 2017; Lissovsky et al. 2017, 2019). Interspecific hybridization and introgression between three species from subgenus *Ochotona* and potential east-west divergence of these species has been hypothesized (Lissovsky et al. 2019, 2022). Population genetic studies in Asian pikas are limited (Zhang et al. 2017; Bhattacharyya and Ishtiaq 2019) and significantly lower than vast literatures on studies of population dynamics and evolution in North American Pikas, for e.g., (Holtz et al. 2016; Robson et al. 2016; Waterhouse et al. 2017; Westover et al. 2020; Klingler et al. 2021).

In this study, we examined populations of five reported species of pikas across their Himalayan and neighboring ranges (Supplementary Table 1, Figure 1) to explore the processes that explains their taxonomic complexity and east-west divergence. We estimated genetic diversity (within and between populations) of six species of pikas within their core and range edges and tested for signals of contemporary and historical gene flow. Based on previous phylogenetic studies, we expect admixture and gene flow among younger and closely related species like *O. curzoniae* and *O. nubrica* (Lissovsky 2014). Alternatively, based on the proximity of the distribution range of species from different subgenera, we also expect gene flow among distant species. Ultimately, we hope the results of this study will expand our knowledge of speciation complexities in Asian Pikas.

**Figure 1:**
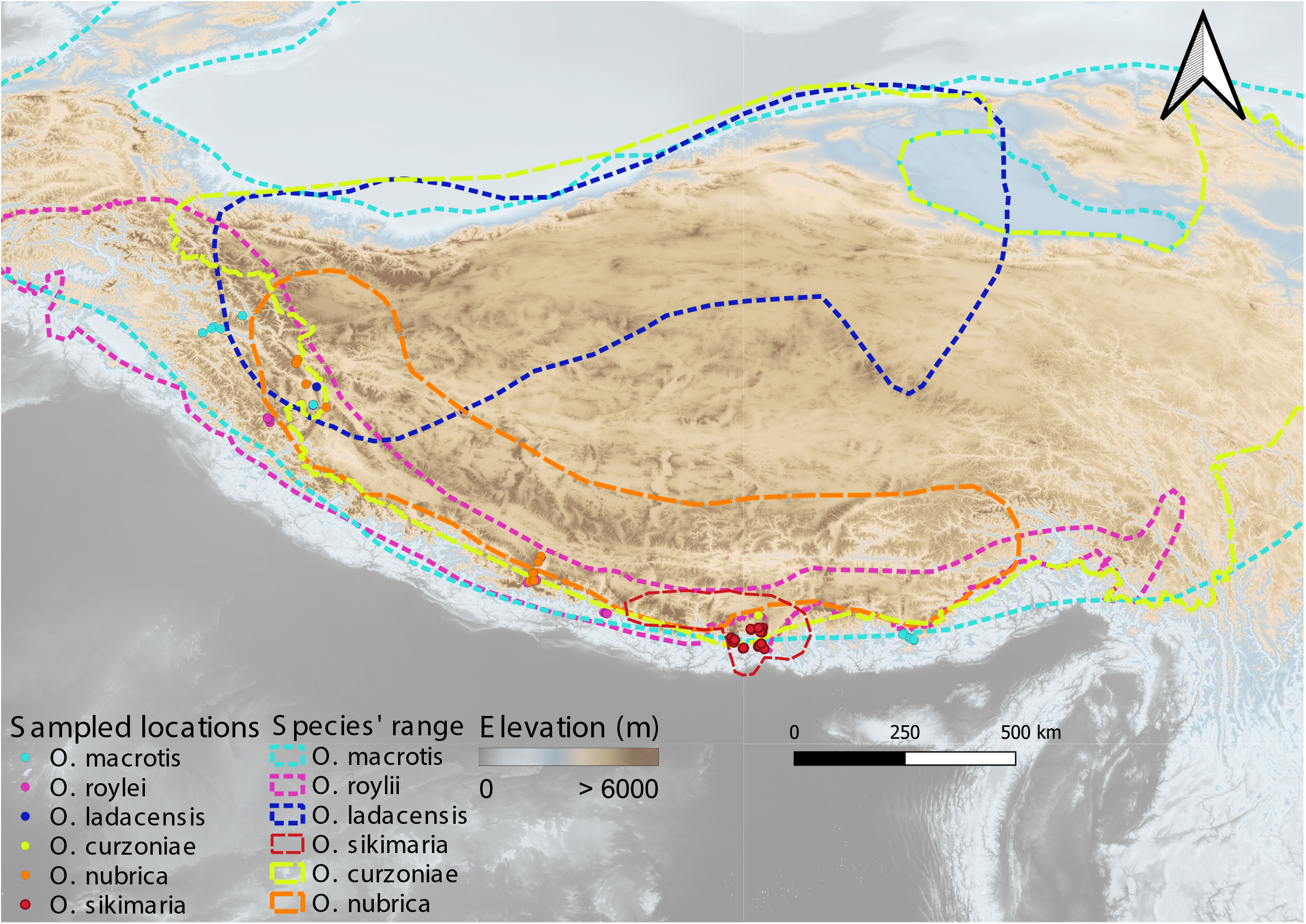
Range map of all six species of Asian pikas across Himalayas and Tibetan Plateau; figure modified from (Dahal et al. 2020). Locations of species sampled are marked by colored dots.

**Figure 2:**
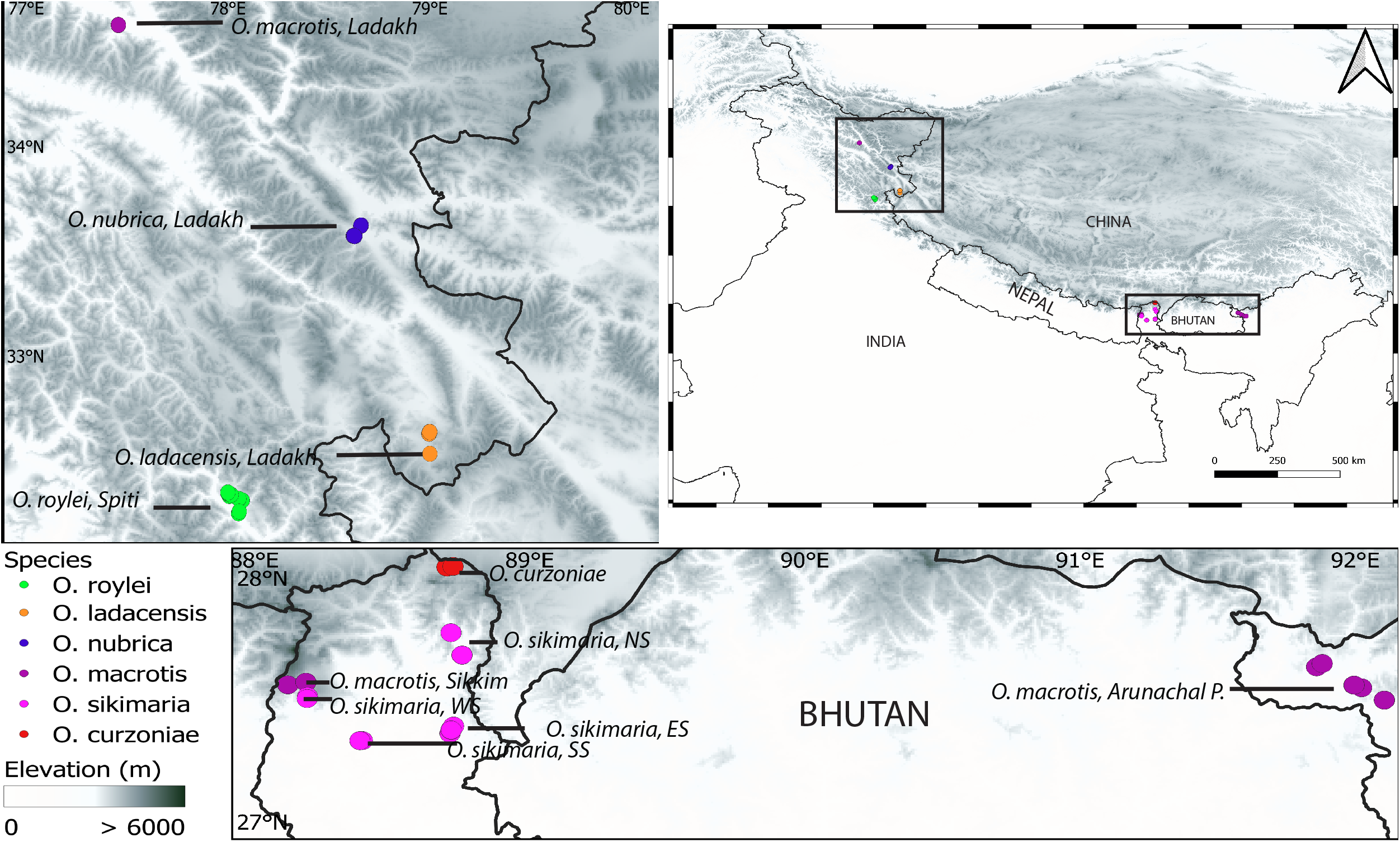
Sampling map. Locations of species sampled in the eastern and western Himalayas are marked with boxes (right panel). Zoomed in sampling map of western (left panel) and eastern (lower panel) Himalayas.

## Methods

### Study area and sample collection

Tissue samples were collected during field surveys carried out in the year 2010 – 2014. Elevational transects were accessed via hiking and traps were set in locations where active pika presence was perceived based on fecal encounter rate and sighting frequencies. Seven sites in eastern Himalaya, five sites in central Himalaya, and seven sites in western Himalaya were selected for trapping pikas and sampling (Supplementary Table 1, Figure 2). In each of these locations, trapping was done for a minimum of three days and trapped individuals were immediately released after taking body measurements and ear punch tissues. Tissue samples were collected in 95%–100% ethanol and stored at −20°C after reaching the laboratory.

### RAD- seq library preparation and sequencing

Genomic DNA was extracted from ear punch tissue samples using Qiagen’s DNeasy Blood and Tissue kit using the manufacturer’s protocol. The double-digest restriction site-associated DNA sequencing (ddRADSeq) library was prepared following the protocol described in (Peterson et al. 2012). The libraries were sequenced at the Centre for Cellular and Molecular Platforms (C-CAMP), an initiative supported by Department of Biotechnology, Govt of India using the Illumina HiSeq platform to generate 100 bp paired-end reads.

### RAD sequences processing

We used STACKS v.2.62 (Catchen et al. 2011) to process the raw sequencing data generated above. Raw sequences were first demultiplexed, barcodes removed, and low-quality sequences were filtered using “process_radtags”. We then mapped paired end “clean” reads from each sample against the *O. curzoniae* genome downloaded from the NCBI database (Sayers et al. 2022). We aligned short sequence reads to the reference genome using bwa v.0.7.17 (Li and Durbin 2009) and further converted the alignment into binary format (bam), sorted the alignment, and removed PCR duplicates using Picard v.2.27.5 (http://broadinstitute.github.io/picard/). We then used the “gstacks” program in STACKS to identify SNPs within the metapopulation for each locus and then genotyped individual at each identified SNP (Catchen et al. 2013) We further used the STACKS “populations” program to analyze a population of individual samples and generated an SNP database in VCF format.

The raw variants were then filtered using a variety of parameters as recommended by standard STACKS workflow practices (https://github.com/enormandeau/stacksworkflow). We only selected genotypes with minimum allele coverage = 2, minimum percent of genotype data per population = 40, the maximum number of populations that can fail percent_genotypes = 3, and the minimum number of samples with rare allele = 2. We then created graphs to find samples with high missing data and removed samples with high missingness (>50% missing genotypes). We further classified SNPs into various categories (singleton, duplicated, diverged, high coverage, low confidence) using the utility scripts provided in STACKS and only kept “singleton” SNPs for downstream analysis. We then estimated pairwise linkage disequilibrium (LD) for each SNP pair using Plink v. 1.09 (Chang et al. 2015) and carried out LD-based pruning of SNPs to keep only one SNP per linked group within the genomic region and make sure there are no spurious correlations among the measured variables. For LD-based pruning, we defined a genomic window of 50kb with a 10bp window step size and prune any SNPs with r^2^ (the measure of LD) > 0.5. Finally, we imputed missing data in the VCF using admixture ancestry relationships (Alexander et al. 2009). The final “clean” SNP dataset consisting of 23,851 variants and 89 samples was used for downstream population genetics analysis.

### Estimation of intra- and inter-species genetic diversity

We used vcftools v.0.1.16 (Danecek et al. 2011) to calculate nucleotide diversity (pi) separately for each species/population using the “cleaned” vcf file. We also calculated the inbreeding coefficient (F) as a measure of relative heterozygosity for each sample using a method of moments implemented in vcftools. The measure of inter-species genetic diversity was calculated using the Fixation index (F_ST_) (Holsinger and Weir 2009).

### Phylogenetic analysis

We examined the genetic relationships among the species by constructing a maximum-likelihood phylogenetic tree using FastTree (Price et al. 2010) with standard parameters for nucleotide alignments of variable positions in the data set. The local support values for each branch in the tree were calculated using Shimodaira-Hasegawa test implemented in FastTree. Timetree database (Kumar et al. 2017) was used to extract the currently known phylogeny of various species of pikas and their closely related species.

### Analysis of population and individual ancestries

We used Admixture v.1.3.0 (Alexander et al. 2009) to estimate individual ancestries from the genome-wide SNP dataset using a maximum-likelihood approach. We first explored the optimal number of genetically distinct clusters that best describes the data using a cross-validation procedure using the K-means method implemented in Admixture. K=7 which had the lowest cross-validation error was chosen as the optimal fit to describe the genetically distinct clusters among the species being studied. We then ran Admixture to assign a cluster to each individual and calculated the proportion of ancestry to the respective cluster. We further plotted the proportion of ancestry data across all samples.

### Exploring evidence of possible gene flow among species

We performed three different analyses to examine gene flow among species. First, we used Treemix v.1.13 (Pickrell and Pritchard 2012) which quantifies the allele frequencies of each SNP to build a maximum-likelihood phylogenetic tree for all defined populations/species and inferrs possible admixture events among the branches. We used 500 bootstrap replicates of the SNP data to test for 10 different migration events (10 replicates for each migration event) and choose an optimum number of migration events. The results indicated that migration event of either 1 or 2 appears optimum (Supplementary Figure 2). We then carried out two runs of Treemix with migration events 1 and 2 to build a consensus tree and identify possible admixture events. To improve the statistical inference, we examined the model residuals to investigate if any parts of the phylogenetic tree are not well modeled for the different runs.

Second, we calculated Patterson’s D-statistics using ABBA-BABA tests (Green et al. 2010; Durand et al. 2011) to examine hybridization and possible gene flow among species. In this test, the number of ancestral (“A”) and derived (“B”) alleles are calculated for a four-taxon comparison that includes an outgroup, predicting patterns termed “ABBA” and “BABA”. Under the condition of incomplete-linage sorting without gene flow, the number of SNPs showing pattern “ABBA” and “BABA” should be equally frequent, whereas an excess of either pattern in the genome indicates possible gene flow between taxa that can be calculated using Patterson’s D-statistics. We used the Dsuite (Malinsky et al. 2021) for calculating D (ABBA/BABA) and f4-ratio statistics for all trios of species in the dataset. As we did not have an outgroup in this dataset, we used two separate internal outgroups (using a different subgenus) based on the phylogenetic tree generated above, (1) *O. nubrica* (subgenus *Ochotona*) to study the pattern of gene flow between trios of *O. roylii, O. macrotis, and O. ladacensis* (subgenus *Conothoa*) (2) *O. roylii* (subgenus *Conothoa*) to study the pattern of gene flow between trios of *O. curzoniae, O. nubrica, and O. sikimaria* (subgenus *Ochotona*).

Third, we inferred pairwise relatedness for each pair of species/population using kinship coefficient using VCFTools.

## Results

### Intra- and inter-species genetic diversity

We used the final filtered SNP dataset consisting of 23,851 variants and 89 samples to calculate the inbreeding coefficient (F) as a measure of observed heterozygosity across all samples. The inbreeding coefficient was generally high across samples and species (F > 0.5) (Supplementary Table 2). Average genome-wide nucleotide diversity was also low across all species/populations (Table 1), consistent with a high inbreeding coefficient.

**Table 1:** Average genome-wide nucleotide diversity across species/populations (the sampling location are shown in parenthesis)

| Species/populations | Nucleotide diversity ($\times 10^{-3}$) |
| --- | --- |
| *O. curzoniae* (North Sikkim) | 0.028 |
| *O. ladacensis* (Ladakh) | 0.020 |
| *O. macrotis* (Arunachal Pradesh) | 0.024 |
| *O. macrotis* (Ladakh) | 0.017 |
| *O. macrotis* (West Sikkim) | 0.033 |
| *O. nubrica* (Ladakh) | 0.020 |
| *O. roylei* (Spiti) | 0.025 |
| *O. sikimaria* (East Sikkim) | 0.022 |
| *O. sikimaria* (North Sikkim) | 0.028 |
| *O. sikimaria* (West and South Sikkim) | 0.030 |

Pairwise-genetic distance (F_ST_) between species ranged from 0.24 to 0.47 and was highest between *O. macrotis* (AP) and *O. sikimaria* (Supplementary Table 3). Intra-species F_ST_ among different populations ranged from 0.03 to 0.28. The lowest pairwise F_ST_ was estimated between *O. macrotis* populations from Ladakh and West Sikkim.

### Characterization of the population structure

We first used the TimeTree database (Kumar et al. 2017) to generate an expected phylogenetic tree based on previously published literature (Supplementary Figure 1). It is expected that *O. nubrica* and *O. curzoniae* being sister species form a phylogenetic cluster and *O. roylei/O. macrotis/O. ladacensis* form a separate cluster. *O. sikimaria* is a newly described species (Dahal et al. 2017) hence is not currently available in the TimeTree database. We further utilized the SNP data to generate a maximum-likelihood phylogenetic tree (Figure 3). All 89 samples were clustered into their respective species/populations. As expected, we observed two major phylogenetic clades (1) consisting of *O. macrotis, O. ladacensis, and O. roylei* (2) consisting of *O. sikimaria, O. nubrica, and O. curzoniae.* Interestingly, few samples from *O. curzoniae* appear intermediate to both clades.

**Figure 3:**
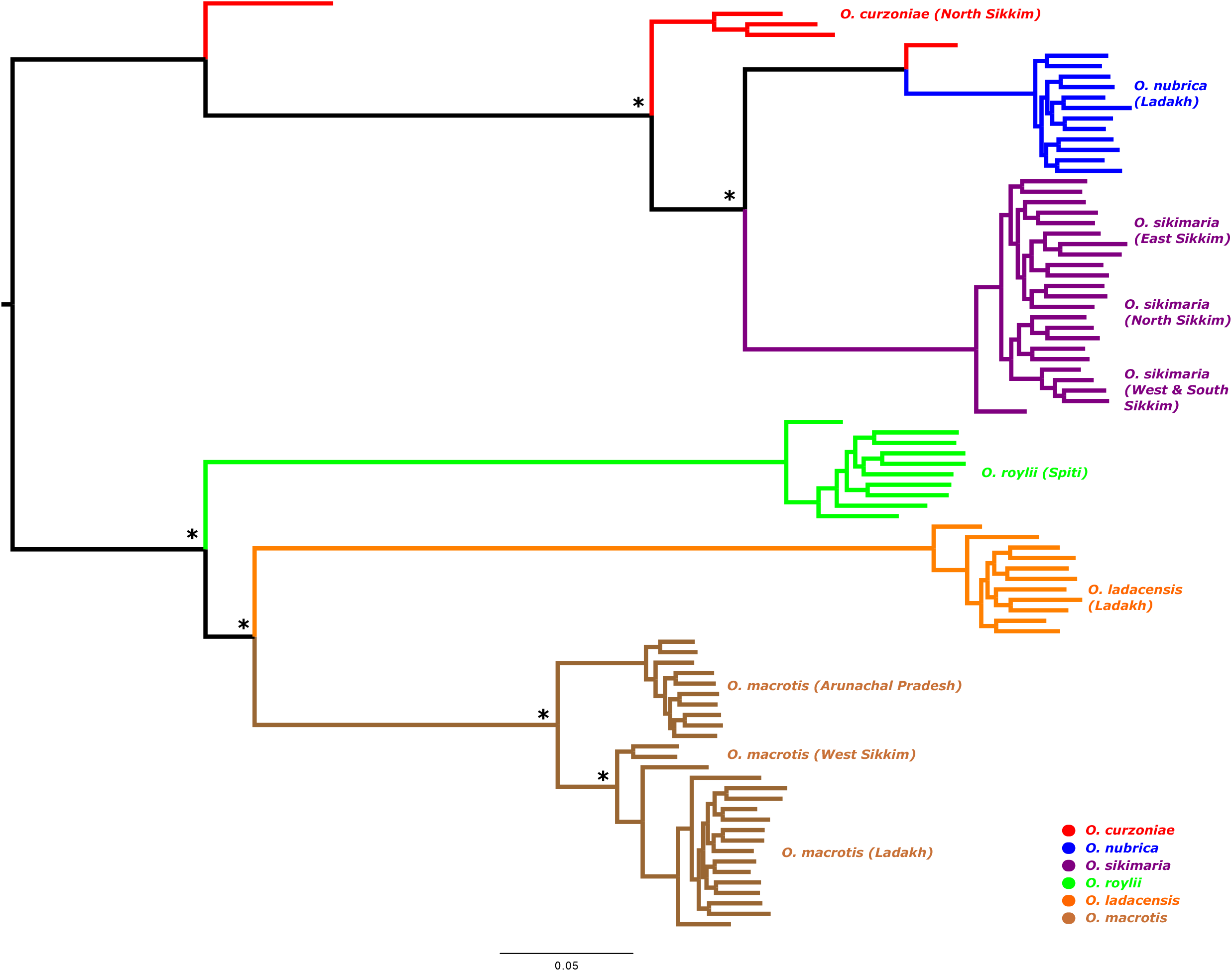
Maximum-likelihood phylogenetic tree based on 23,851 genome-wide SNPs and 89 samples from six species of Asian pikas. All nodes having full local support based on the Shimodaira–Hasegawa test (Price et al. 2010) are marked by asterisks.

We also inferred the population and individual ancestries of all samples using the K-means method using Admixture (Alexander et al. 2009). A cross-validation procedure indicated K=7 as the optimal number of the genetic cluster (Supplementary Table 4). The proportion of ancestry for K=7 (Figure 4) was overall consistent with the phylogenetic tree (Figure 3). We found the *O. macrotis* populations from Ladakh (LK) and West Sikkim (WS) are similar, whereas the Arunachal population (AP) appears genetically distinct. We further calculated pairwise average genetic distance among these three populations of *O. macrotis* (Table 2). The population from AP is 11.24% divergent from LK and 8.78% divergent from WS, indicating its genetic distinctness.

**Figure 4:**
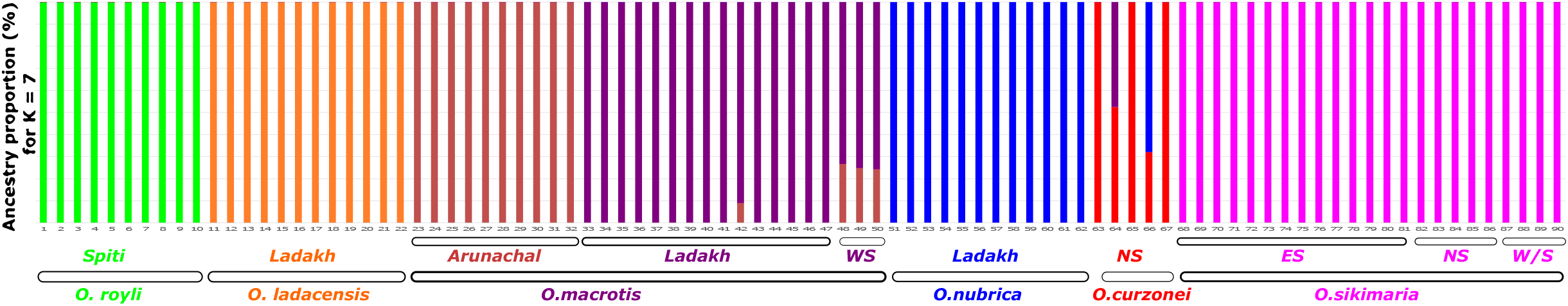
Admixture plot illustrating ancestry among Asian pikas for K=7. Individuals are shown as vertical bars colored in proportion to their estimated ancestry within each cluster.

**Table 2:** Average pairwise-genetic distance among three populations of O. macrotis (AP: Arunachal Pradesh, WS: West Sikkim and LK: Ladakh)

|  | AP | LK | WS |
| --- | --- | --- | --- |
| AP | 3.66 | 11.24 | 8.78 |
| LK | 9.87 | 5.58 | 7.35 |
| WS | 8.78 | 7.35 | 3.66 |

All samples of *O. sikimaria* from different locations in Sikkim as well as *O. roylii* and *O. ladacensis* are genetically distinct (Figure 4). Some samples of *O. curzoniae* appear to share ancestry with *O. nubrica* and *O. macrotis* samples from Ladakh.

### Possible gene flow among species

Results from the phylogenetic tree and inference of population and individual ancestries indicated possible gene sharing among some of these species. Hence, we performed three different analyses (1) Treemix (Pickrell and Pritchard 2012) (2) ABBA-BABA (Green et al. 2010; Durand et al. 2011), and (3) Genetic relatedness (Manichaikul et al. 2010) to examine the possible evidence of gene sharing among different species. We first ran TreeMix with bootstrapping, which helped us to choose an optimum number of migration events in our dataset and create a consensus tree. We plotted the treemix plot with migration events 1 and 2 (Figure 5). If we consider one migration event, our data indicated gene flow between *O. curzoniae* and *O. ladacensis.* If we consider two migration events, the data indicates (a) the first gene flow between *O. curzoniae* and the base of *O. ladacensis/ O. macrotis* split and (b) between Ladakh populations of *O. ladacensis* and *O. macrotis.*

**Figure 5:**
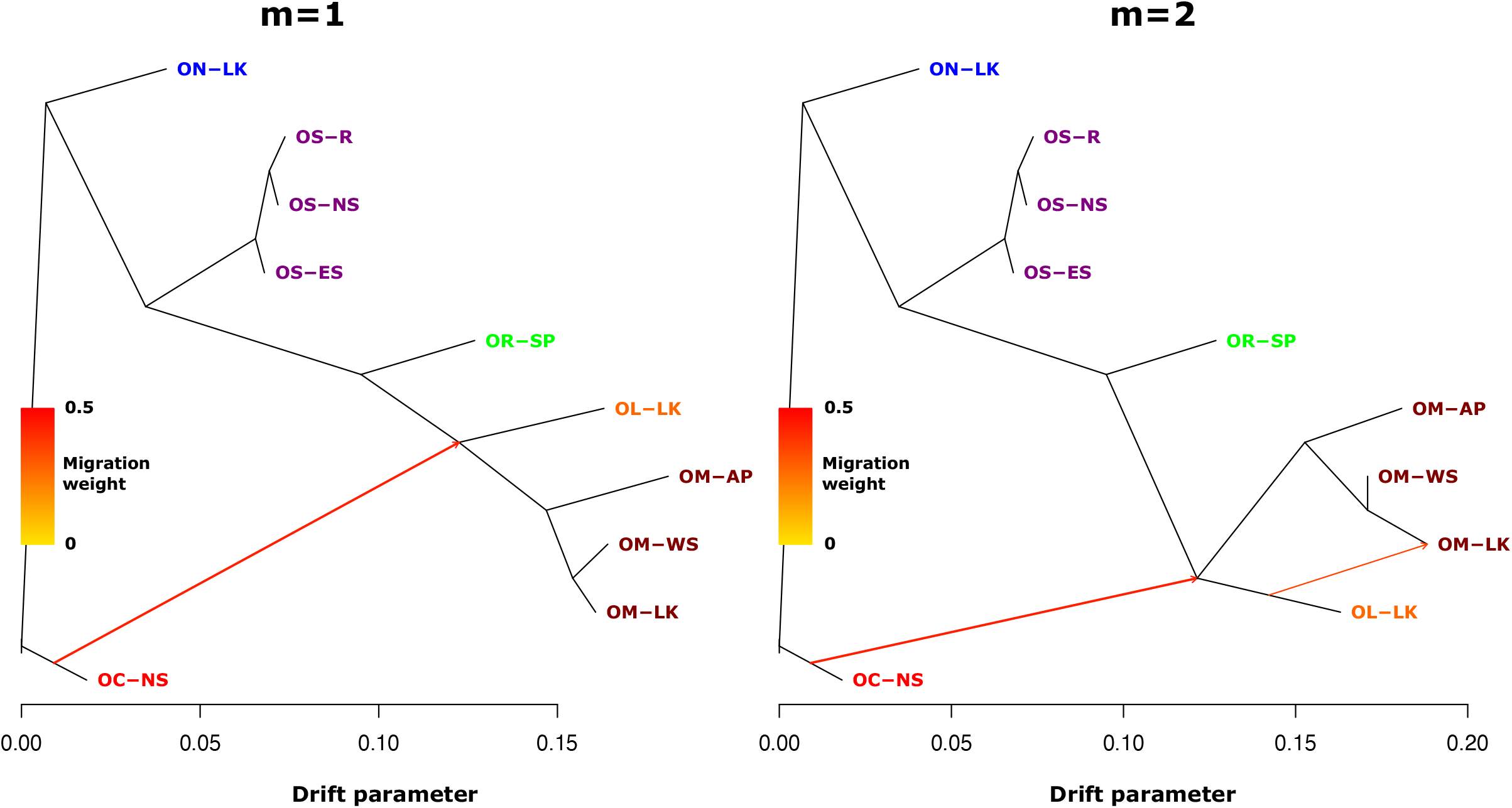
Population graphs inferred by TreeMix (Pickrell and Pritchard 2012) using 23,851 genome-wide SNPs. Branch lengths are proportional to the evolutionary change (the drift parameter) and terminal nodes are labeled with population codes (see Supplementary Table 1 for details). TreeMix phylogram is shown with one (left panel) or two migration events (right panel).

We further examined the pattern of gene flow between species using D and f4-ratio statistics, based on studying correlations of allele frequencies across populations implemented in Dsuite (Malinsky et al. 2021). As we did not have an outgroup in this dataset, we used two separate internal outgroups based on the phylogenetic tree generated above, (1) *O. nubrica* to study the pattern of gene flow between trios of *O. roylii, O. macrotis, and O. ladacensis* (2) *O. roylii* to study the pattern of gene flow between trios of *O. curzoniae, O. nubrica, and O. sikimaria*. For scenario 1, we found evidence of excess allele sharing between populations of *O. macrotis* from AP and WS, but not between AP and LK (Supplementary Figure 3). We also found evidence of gene flow between Ladakh populations of *O. ladacensis* and *O. macrotis.* For scenario 2, we found evidence of gene flow between populations of *O. sikimaria* and *O. nubrica* (Supplementary Figure 3).

We also calculated relatedness statistics based on the method of (Manichaikul et al. 2010) for each pair of individuals. The results are consistent with the Treemix and ABBA-BABA analysis. *O. curzoniae* shows a higher degree of relatedness with *O. macrotis* and *O. nubrica* (Figure 6). *O. macrotis* populations from Ladakh and West Sikkim are similar, and the Arunachal population appears distinct.

**Figure 6:**
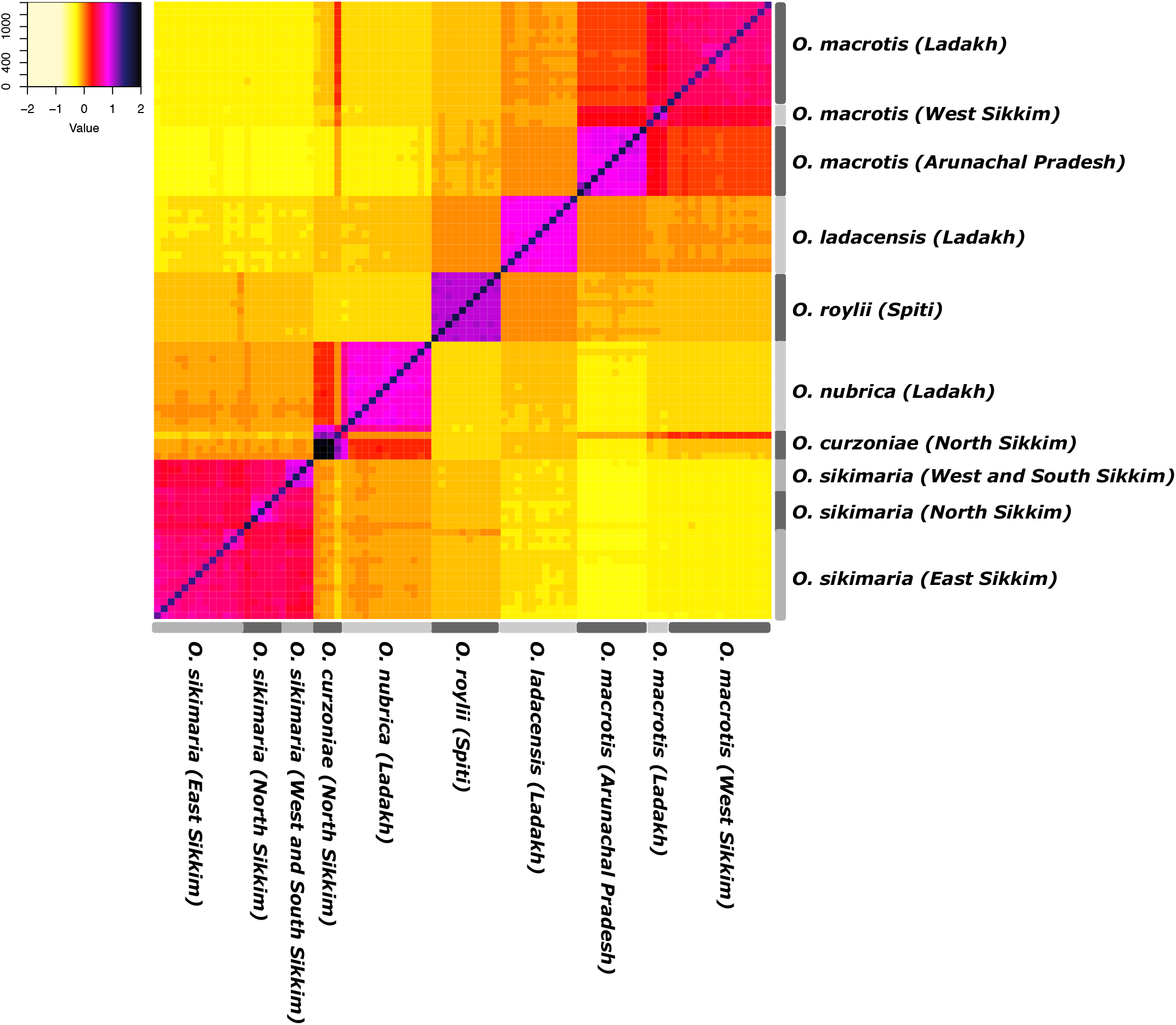
Heatmap showing relatedness score (Manichaikul et al. 2010) for each pair of individuals. Higher values indicate individuals share higher genetic similarities.

## Discussion

Here, we study six species of Pikas (*Ochotona* spp.) that are distributed across the eastern and western Himalayas and neighboring high-altitude mountain regions (Figure 1, 2 and Supplementary Table 1). This sampling included three populations of *O. macrotis* distributed across the eastern and western Himalayas. We utilized a genomic approach using more than 20,000 genome-wide SNP markers obtained from double digest restriction-site associated DNA sequencing (ddRAD-seq) to estimate intra- and inter-species genetic diversity, population structure, and phylogenetic history and explore processes that shaped the current genetic diversity of Pikas across the Himalayas.

### Genetic diversity estimates indicate decreasing population trend in Asian pikas

The nucleotide diversity was consistently low across all species (Table 2) whereas the inbreeding coefficient was high (>0.5) (Supplementary Table 2). These two estimates indicated that all six species of pikas in this study had low genetic diversity. A previous study in *O. curzoniae* using 10 microsatellite markers had also identified a high inbreeding coefficient in this species (Zhang et al. 2017), which is consistent with our findings based on a larger set of genome-wide markers. Low genetic diversity is also a common feature in American pikas (*O. princeps*) (Robson et al. 2016; Klingler et al. 2021), which has been associated with elevation-specific population decline and restricted dispersal. Low genetic diversity in Asian pikas may also be an indication of decreasing population size. In addition, the Himalayas are the range edge for most of these species, especially *O. curzoniae* and *O. ladacensis* (Dahal et al. 2020) which may also contribute to low population size and thereby, low nucleotide diversity. However, to investigate the edge effect on the genetic diversity of Asian Pikas, we need data from across their elevation limits, including their core range for wide-ranging species like *O. macrotis* and *O. roylei.*

### Is the genetic distance between species explained by geographic distance?

Inter-species pairwise genetic differentiation (F_ST_) was generally high for each species pair, and most were consistent with the geographic distance between species (Supplementary Table 3). But there were certain cases where inter-species pairwise genetic differentiation did not correspond to the geographic distance between species. For e.g., the population of *O. macrotis* from Sikkim showed strong genetic differentiation (F_ST_ = 0.28) with the population from Arunachal Pradesh (both eastern Himalayas) whereas very low genetic differentiation with the population from Ladakh (western Himalayas). Higher divergence in population of *O. macrotis* from Arunachal Pradesh is consistent with the previous study that indicated *O. macrotis* from Arunachal Pradesh may be as a different subspecies (Lissovsky et al. 2017). Low divergence among geographically distant populations of *O. macrotis* in the eastern and western Himalayas indicated the possibility of historic gene flow among ancestral populations of this species.

### Gene flow between species

We further explored population structure and patterns of gene flow to examine the processes associated with the intra- and inter-species genetic diversity. The phylogenetic tree based on SNP data (Figure 3) was consistent with previously known phylogeny (Supplementary Figure 1). *O. macrotis, O. ladacensis, and O. roylei* were grouped into one cluster which is consistent as these species are from the same subgenus (*Conothoa*) (Supplementary Table 1). Similarly, *O. sikimaria, O. nubrica, and O. curzoniae* were grouped into another cluster which is also consistent as these species are from different subgenus (*Ochotona*). Interestingly, few samples from *O. curzoniae* appear intermediate to both clades which is a possible indication of gene flow across species. Cases of interspecific hybridization and introgression among three species of subgenus Ochotona has been previously hypothesized (Lissovsky et al. 2019).

The inference of population ancestry (Figure 4) was consistent with the population structure identified based on phylogenetic analysis. Some individuals *O. curzoniae* appeared to have mixed ancestry which was consistent with their relatively intermediate position in the phylogenetic tree. *O. macrotis* populations in West Sikkim shared the majority of ancestry with those from Ladakh. It is consistent with low genetic divergence (F_ST_) we have identified among these two populations (Supplementary Table 3). The genetic similarity of these two populations is interesting considering their geographic isolation (one from eastern and other from western Himalayas) (Figures 1 and 2). However, *O. macrotis* has a wider distribution range (Figure 1), and this observed genetic similarity possibly indicates patterns of historic gene flow in their ancestral populations. The admixture ancestry showed the *O. macrotis* population from Arunachal Pradesh as genetically distinct, which supports the previous classification of this population as a separate subspecies (Lissovsky et al. 2017).

Treemix results indicated possible gene flow between *O. curzoniae* and *O. ladacensis.* The distribution range of these two species and the populations included in the current study overlap at their range edge (Figure 1) (Dahal et al. 2020) and have similar species ecology (Table 1), hence these species have a possibility of sharing genes. On the contrary, gene flow between non-sibling species, *O. curzoniae,* and *O. ladacensis* were unexpected, but such patterns are not unique to the current study (Stölting et al. 2013). We also identified evidence of gene flow among populations of *O. ladacensis* and *O. macrotis* in Ladakh (eastern Himalayas). ABBA-BABA analysis indicated there was excess allele sharing (that cannot be explained by incomplete lineage sorting) among populations of *O. macrotis* in the eastern Himalayas, but not between populations between the eastern and western Himalayas. Consistent with Treemix, this analysis also independently identified evidence of gene flow among populations of *O. ladacensis* and *O. macrotis* in Ladakh. It also identified evidence of gene flow between populations of *O. sikimaria* and *O. nubrica*. This is consistent with previously hypothesized cases of interspecific hybridization and introgression among these species (Lissovsky et al. 2019). Finally, the estimates of pairwise relatedness scores among species/populations were also consistent with the results of Treemix and ABBA-BABA analysis (Figure 6). Identification of similar patterns of gene flow among species using three separate analytical methods indicated that processes of historical gene flow across the phylogeny have been a key feature during the evolution of Asian pikas.

Our results have identified two important processes associated with the divergence and speciation of Himalayan Pikas (a) low genetic diversity and (b) extensive evidence of gene flow across species. Genetic diversity is a major contributor to the evolutionary viability of natural populations and has a key implication in species conservation (Frankham et al., 2010). Low genetic diversity indicates that the effective population size of Asian pikas is possibly declining. These results are consistent with species distribution models predicted for future habitat loss in certain species e.g. *O. roylei* (Bhattacharyya et al. 2019). Although Asian Pikas currently have “Least Concern” status on IUCN red list (IUCN 2022), our results indicate that the need for effective management and monitoring practices are being essential. However, it appears that inter-species gene flow that introduces new alleles into the population has been a key evolutionary process among Asian Pikas to counter the negative effect of low genetic diversity. The current study shows that speciation with gene flow is a common pattern in Asian pikas. However, high-quality reference genomes from each of these species and population-scale whole-genome sequencing will be required in future to better characterize genomic signatures of species hybridization and gene flow.

## Data accessibility

All genomic data generated in this study is available via NCBI BioProject, accession number PRJNA893189

(https://www.ncbi.nlm.nih.gov/bioproject/PRJNA893189).

## Acknowledgments

We acknowledge field assistants Sukhraj, B.L Subba, Sunil Subba, and the team from East Sikkim, Bhupendra Dahal, Raju, Dorjay, Rinchen, Sherap, and Rigzin Dorjay for their help in the fieldwork work and sampling. We also acknowledge the forest departments of Arunachal Pradesh, Sikkim (especially Range Officer, CC Lachungpa and his team), Himachal Pradesh, and Ladakh (especially Mr. Intesar Suhel) for providing necessary collection permit and logistic support. We also thank the Centre for Cellular and Molecular Platforms (C-CAMP’s) Next Generation Genomics facility for carrying out the ddRAD sequencing run. This work was supported by the Department of Science and Technology (DST), Govt. of India for the DST INSPIRE Faculty Award (grant number DST/INSPIRE/04/2018/001587) to ND, Sikkim Department of Biotechnology, Government of India, project to URK and faculty start-up funds (grant number 201113) from Kent State University, Department of Biological Sciences to SL.

## Author contributions

ND, UR, and SL conceived and designed the study. ND conducted the field and lab work. SK contributed samples from East Sikkim. ND, MR, and SL analyzed the data. ND and SL wrote the manuscript; UR and SL edited the manuscript and interpreted the results.

