## Supplementary Materials for "Genomic Variation, Population Structure, and Gene Flow across Asian Pikas"

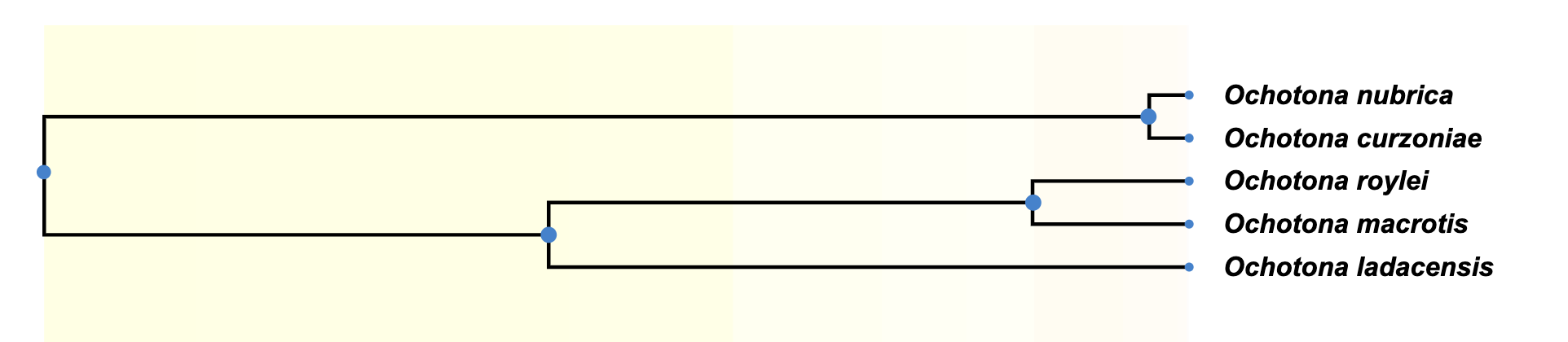

***Supplementary Fig 1:*** *Expected phylogenetic tree based on Timetree database (Kumar et al. 2017) for all species of pikas currently used in this study. O. sikimaria is a newly described species* *(Dahal et al. 2017) hence is not currently available in the TimeTree database.*

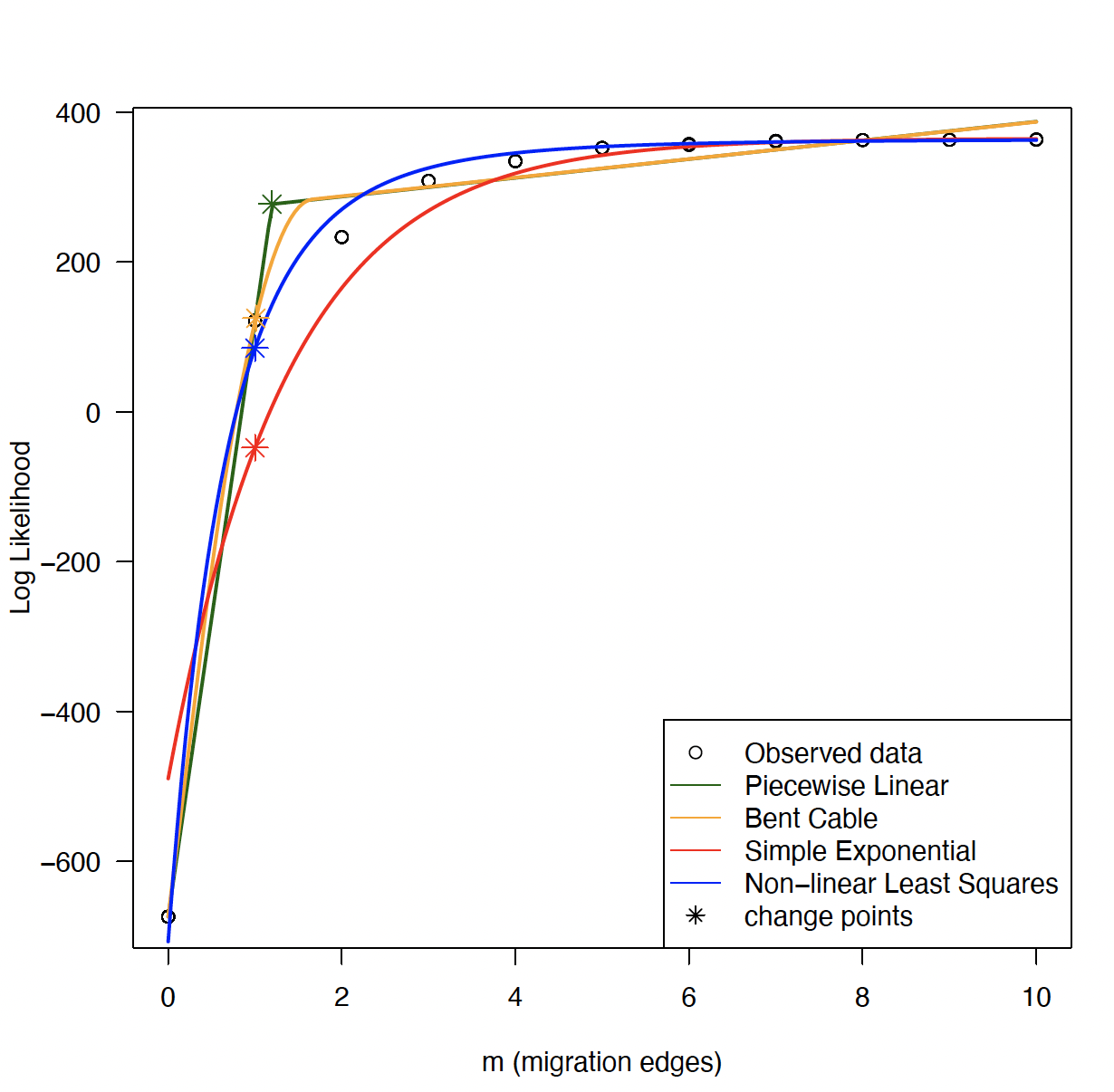

***Supplementary Figure 2:*** *Plot showing changes in log-likelihood for a possible number of migration events (m). Migration event of either 1 or 2 appears optimum.*

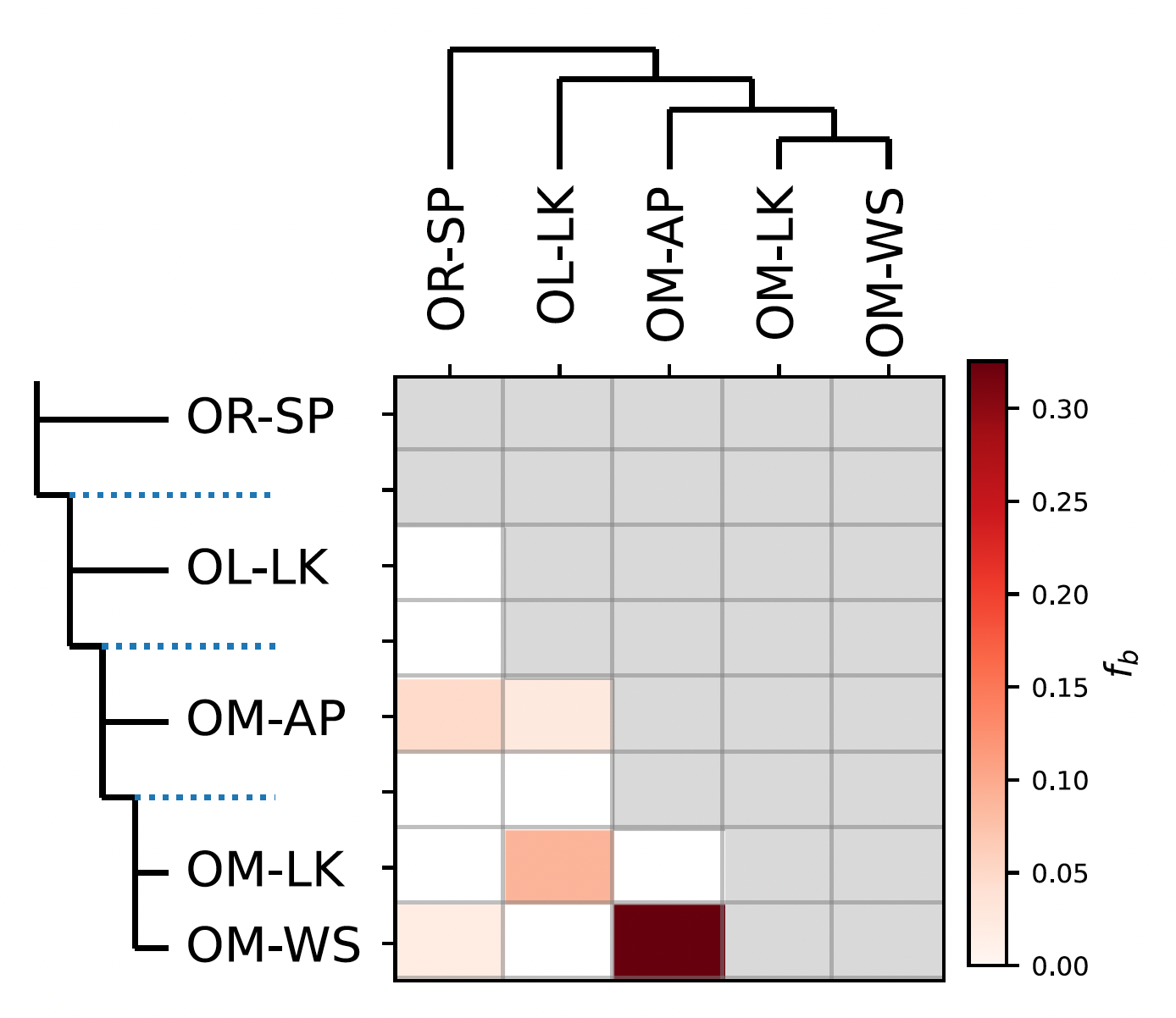

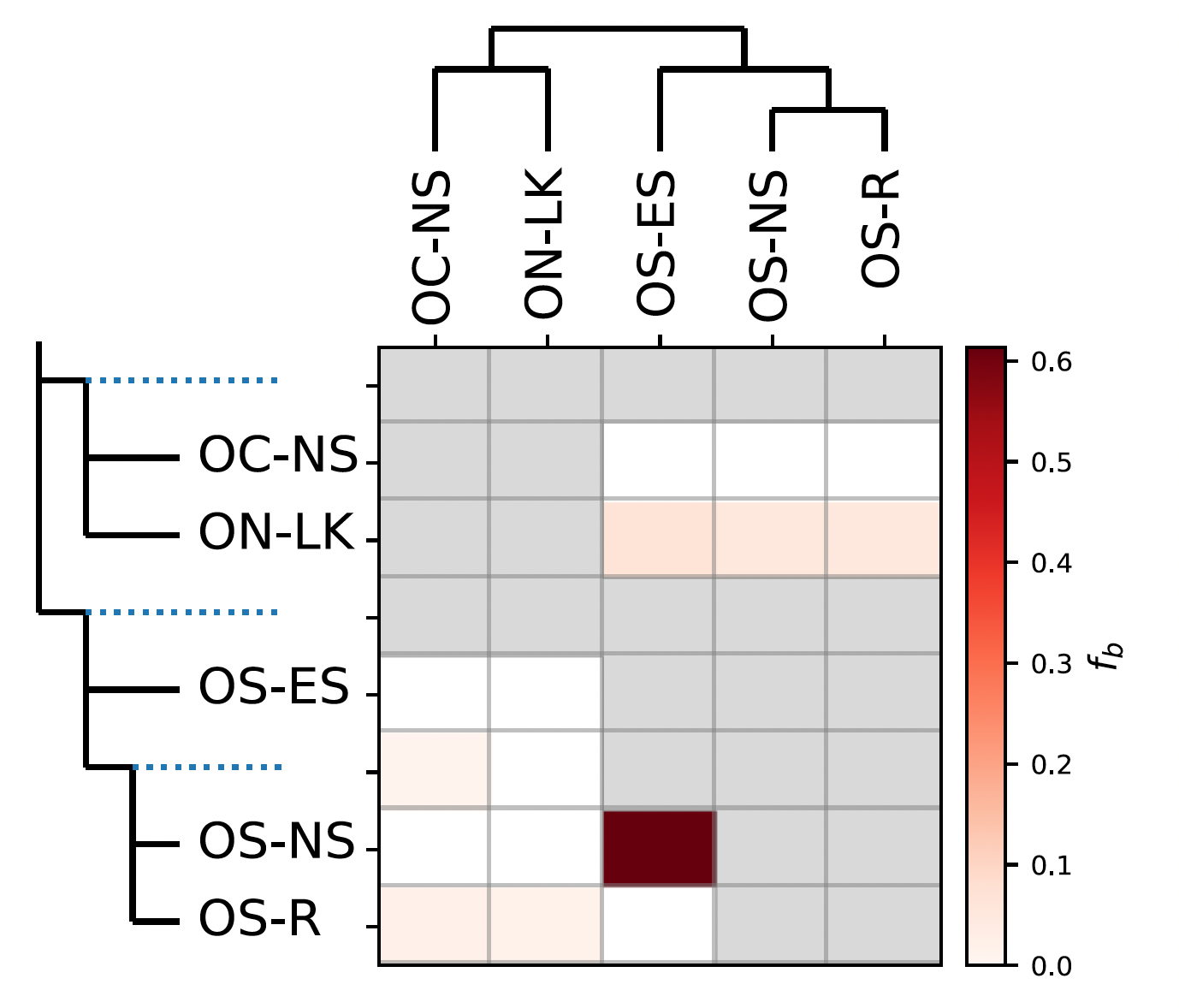

***Supplementary Figure 3:*** *Heatmap summarizing the f-branch statistics estimated by Dsuite* *(Malinsky et al., 2021)**. Darker colors depict increasing evidence for gene flow between lineages. Dotted lines in the phylogeny represent ancestral lineages.*

***Supplementary Table 1:*** *Sampling location and brief habitat/ecology of the Asian pika species sampled in this study.*

| **Species (population code)** | **Sampled location** | **Range** | **Sampled elevation (m)** | **IUCN red list elevation range (in m)** | **Subgenera** | **Sampled habitat type** | **Species ecology in brief** |
| --- | --- | --- | --- | --- | --- | --- | --- |
| *O. curzoniae*  (OC-NS) | North Sikkim | Tibetan plateau | 5060 - 5067 | 3000 - 5000 | Ochotona | Open desert - steppe | Burrowing pika, makes complex burrows; social |
| *O. ladacensis*  (OL-LK) | Ladakh | Western Himalaya | 4840-4888 | 4200 - 5400 | Conothoa | Open desert-steppe | Burrowing pika, social |
| *O. macrotis*  (OM-AP) | Arunachal Pradesh | Eastern Himalayas | 3311-4213 | 2300-6400 | Conothoa | Big rocky patches | Talus dweller, asocial |
| *O.macroitis*  (OM-LK) | Ladakh | Eastern and Western Himalayas | 3205-3212 | 2300-6400 | Conothoa | Rocky patches near stream | Talus dweller, asocial |
| *O. macrotis*  (OM-WS) | West Sikkim | Eastern and Western Himalayas | 4559-4754 | 2300-6400 | Conothoa | Big rocky patches near lake | Talus dweller, asocial |
| *O. nubrica*  (ON-LK) | Ladakh | Central to Eastern Himalayas | 4330-4403 | 3000-4500 | Ochotona | Shrub steppe marshes, dominated by *Caragana* sp. | Burrowing pika (shallow burrower), shy |
| *O. roylei*  (OR-SP) | Spiti, Himachal Pradesh | Eastern and Western Himalayas | 3649-4997 | 2400-5200 | Conothoa | Rocky patches | Talus dweller, asocial |
| *O. sikimaria*  (OS-ES) | East Sikkim | Eastern and Western Himalayas | 3096-3666 | not assessed | Ochotona | Sub- alpine Rhododendron Forest | Burrows in open habitat with many entrances; burrows in farmlands are less complex; social |
| *O. sikimaria*  (OS-NS) | North Sikkim | Eastern Himalayas | 2961-3509 | not assessed | Ochotona | Sub- alpine forest: Farmland and Rhododendron forests. | Burrows in open habitat with many entrances; burrows in farmlands are less complex; social |
| *O. sikimaria*  OS-R | South and West Sikkim | Eastern Himalayas | 3233-4203 | not assessed | Ochotona | Sub- alpine forest dominated with shrubs of R*hododendron* sp. and *Gaultheria* sp. | Burrows in open habitat with many entrances; burrows in farmlands are less complex; social |

**Supplementary Table 2**: Inbreeding coefficient of each sample. See Supplementary Table 1 for species and population code.

| **INDIVIDUAL** | **O(HOM)** | **E(HOM)** | **N_SITES** | **F** |
| --- | --- | --- | --- | --- |
| OC-NS_2d18 | 22636 | 19844.8 | 23851 | 0.69672 |
| OC-NS_2d20 | 20768 | 19844.8 | 23851 | 0.23045 |
| OC-NS_2d21b | 22641 | 19844.8 | 23851 | 0.69797 |
| OC-NS_2d22 | 21748 | 19844.8 | 23851 | 0.47507 |
| OC-NS_2d26 | 22592 | 19844.8 | 23851 | 0.68574 |
| OL-LK_L25 | 22670 | 19844.8 | 23851 | 0.70521 |
| OL-LK_L26 | 22693 | 19844.8 | 23851 | 0.71095 |
| OL-LK_L27 | 22605 | 19844.8 | 23851 | 0.68898 |
| OL-LK_L28 | 22722 | 19844.8 | 23851 | 0.71819 |
| OL-LK_L30 | 22676 | 19844.8 | 23851 | 0.70671 |
| OL-LK_L31 | 22710 | 19844.8 | 23851 | 0.71519 |
| OL-LK_L32 | 22749 | 19844.8 | 23851 | 0.72493 |
| OL-LK_L33 | 22759 | 19844.8 | 23851 | 0.72742 |
| OL-LK_L34 | 22632 | 19844.8 | 23851 | 0.69572 |
| OL-LK_L35 | 22645 | 19844.8 | 23851 | 0.69897 |
| OL-LK_L37 | 22743 | 19844.8 | 23851 | 0.72343 |
| OM-AP_7d11 | 23079 | 19844.8 | 23851 | 0.8073 |
| OM-AP_7d3 | 23071 | 19844.8 | 23851 | 0.8053 |
| OM-AP_A7-10 | 23121 | 19844.8 | 23851 | 0.81778 |
| OM-AP_A7-14 | 23011 | 19844.8 | 23851 | 0.79033 |
| OM-AP_A7-31 | 23101 | 19844.8 | 23851 | 0.81279 |
| OM-AP_A7-32 | 22933 | 19844.8 | 23851 | 0.77086 |
| OM-AP_A7-33 | 22953 | 19844.8 | 23851 | 0.77585 |
| OM-AP_A7-4 | 23088 | 19844.8 | 23851 | 0.80955 |
| OM-AP_A7-7 | 23199 | 19844.8 | 23851 | 0.83725 |
| OM-AP_A7-8 | 23089 | 19844.8 | 23851 | 0.8098 |
| OM-LK_A29 | 22866 | 19844.8 | 23851 | 0.75413 |
| OM-LK_A32 | 22947 | 19844.8 | 23851 | 0.77435 |
| OM-LK_L20 | 22805 | 19844.8 | 23851 | 0.73891 |
| OM-LK_L23 | 22948 | 19844.8 | 23851 | 0.7746 |
| OM-LK_L38 | 22802 | 19844.8 | 23851 | 0.73816 |
| OM-LK_L39 | 22753 | 19844.8 | 23851 | 0.72593 |
| OM-LK_L40 | 22671 | 19844.8 | 23851 | 0.70546 |
| OM-LK_L42 | 22646 | 19844.8 | 23851 | 0.69922 |
| OM-LK_L43 | 22874 | 19844.8 | 23851 | 0.75613 |
| OM-LK_L44 | 22616 | 19844.8 | 23851 | 0.69173 |
| OM-LK_L45 | 22689 | 19844.8 | 23851 | 0.70995 |
| OM-LK_L46 | 22810 | 19844.8 | 23851 | 0.74015 |
| OM-LK_L47 | 22791 | 19844.8 | 23851 | 0.73541 |
| OM-LK_L48 | 22673 | 19844.8 | 23851 | 0.70596 |
| OM-LK_LqH | 22628 | 19844.8 | 23851 | 0.69473 |
| OM-WS_5d4 | 22863 | 19844.8 | 23851 | 0.75338 |
| OM-WS_5d8 | 22851 | 19844.8 | 23851 | 0.75039 |
| OM-WS_5d9 | 22456 | 19844.8 | 23851 | 0.65179 |
| ON-LK_L1 | 22698 | 19844.8 | 23851 | 0.7122 |
| ON-LK_L10 | 22790 | 19844.8 | 23851 | 0.73516 |
| ON-LK_L14 | 22823 | 19844.8 | 23851 | 0.7434 |
| ON-LK_L15 | 22650 | 19844.8 | 23851 | 0.70022 |
| ON-LK_L16 | 22692 | 19844.8 | 23851 | 0.7107 |
| ON-LK_L18 | 22588 | 19844.8 | 23851 | 0.68474 |
| ON-LK_L29 | 22663 | 19844.8 | 23851 | 0.70346 |
| ON-LK_L4 | 22750 | 19844.8 | 23851 | 0.72518 |
| ON-LK_L6 | 22736 | 19844.8 | 23851 | 0.72168 |
| ON-LK_L8 | 22669 | 19844.8 | 23851 | 0.70496 |
| ON-LK_L9 | 22665 | 19844.8 | 23851 | 0.70396 |
| ON-LK_LqCHU | 22609 | 19844.8 | 23851 | 0.68998 |
| OR-SP_4d1 | 22471 | 19844.8 | 23851 | 0.65554 |
| OR-SP_4d11 | 22368 | 19844.8 | 23851 | 0.62983 |
| OR-SP_4d12 | 22418 | 19844.8 | 23851 | 0.64231 |
| OR-SP_4d14 | 22427 | 19844.8 | 23851 | 0.64455 |
| OR-SP_4d16 | 22413 | 19844.8 | 23851 | 0.64106 |
| OR-SP_4d27 | 22453 | 19844.8 | 23851 | 0.65104 |
| OR-SP_4d3 | 22404 | 19844.8 | 23851 | 0.63881 |
| OR-SP_4d9 | 22384 | 19844.8 | 23851 | 0.63382 |
| OR-SP_A30 | 22444 | 19844.8 | 23851 | 0.6488 |
| OR-SP_A31 | 22462 | 19844.8 | 23851 | 0.65329 |
| OS-ES_8d11 | 21368 | 19844.8 | 23851 | 0.38021 |
| OS-ES_8d12 | 21824 | 19844.8 | 23851 | 0.49404 |
| OS-ES_8d13 | 21954 | 19844.8 | 23851 | 0.52649 |
| OS-ES_8d16 | 21883 | 19844.8 | 23851 | 0.50876 |
| OS-ES_8d18 | 21962 | 19844.8 | 23851 | 0.52848 |
| OS-ES_8d2 | 21721 | 19844.8 | 23851 | 0.46833 |
| OS-ES_8d3 | 21887 | 19844.8 | 23851 | 0.50976 |
| OS-ES_8d5 | 21781 | 19844.8 | 23851 | 0.4833 |
| OS-ES_8d8 | 22040 | 19844.8 | 23851 | 0.54795 |
| OS-ES_8d9 | 21852 | 19844.8 | 23851 | 0.50103 |
| OS-ES_9d11 | 21868 | 19844.8 | 23851 | 0.50502 |
| OS-ES_9d13 | 21955 | 19844.8 | 23851 | 0.52674 |
| OS-ES_9d14 | 21914 | 19844.8 | 23851 | 0.5165 |
| OS-ES_9d7 | 21884 | 19844.8 | 23851 | 0.50901 |
| OS-NS_1d1 | 22787 | 19844.8 | 23851 | 0.73441 |
| OS-NS_1d2 | 22182 | 19844.8 | 23851 | 0.5834 |
| OS-NS_1d3 | 22414 | 19844.8 | 23851 | 0.64131 |
| OS-NS_1d6 | 21748 | 19844.8 | 23851 | 0.47507 |
| OS-NS_1d7 | 22172 | 19844.8 | 23851 | 0.5809 |
| OS-R_2d23 | 22547 | 19844.8 | 23851 | 0.67451 |
| OS-R_2d24 | 22613 | 19844.8 | 23851 | 0.69098 |
| OS-R_2d25 | 22604 | 19844.8 | 23851 | 0.68873 |
| OS-R_5d3 | 22882 | 19844.8 | 23851 | 0.75813 |

***Supplementary Table 3:*** *Pairwise FST between species/population. See Supplementary Table 1 for species and population code*

|  | **OC-NS** | **OL-LK** | **OM-AP** | **OM-LK** | **OM-WS** | **ON-LK** | **OR-SP** | **OS-ES** | **OS-NS** | **OS-R** |
| --- | --- | --- | --- | --- | --- | --- | --- | --- | --- | --- |
| **OC-NS** | 0 | 0.33 | 0.44 | 0.31 | 0.34 | 0.27 | 0.38 | 0.24 | 0.25 | 0.26 |
| **OL-LK** | 0.33 | 0 | 0.32 | 0.25 | 0.3 | 0.34 | 0.31 | 0.27 | 0.32 | 0.34 |
| **OM-AP** | 0.44 | 0.32 | 0 | 0.2 | 0.28 | 0.44 | 0.41 | 0.33 | 0.43 | 0.47 |
| **OM-LK** | 0.31 | 0.25 | 0.2 | 0 | 0.03 | 0.31 | 0.3 | 0.26 | 0.31 | 0.33 |
| **OM-WS** | 0.34 | 0.3 | 0.28 | 0.03 | 0 | 0.43 | 0.39 | 0.31 | 0.35 | 0.39 |
| **ON-LK** | 0.27 | 0.34 | 0.44 | 0.31 | 0.43 | 0 | 0.39 | 0.25 | 0.32 | 0.35 |
| **OR-SP** | 0.38 | 0.31 | 0.41 | 0.3 | 0.39 | 0.39 | 0 | 0.27 | 0.33 | 0.36 |
| **OS-ES** | 0.24 | 0.27 | 0.33 | 0.26 | 0.31 | 0.25 | 0.27 | 0 | 0.03 | 0.05 |
| **OS-NS** | 0.25 | 0.32 | 0.43 | 0.31 | 0.35 | 0.32 | 0.33 | 0.03 | 0 | 0.02 |
| **OS-R** | 0.26 | 0.34 | 0.47 | 0.33 | 0.39 | 0.35 | 0.36 | 0.05 | 0.02 | 0 |

***Supplementary Table 4****: Cross-validation error (CV) for k=1-11 for admixture analysis. The lowest error K=7) indicated the optimal number of genetic clusters.*

CV error (K=1): 0.85100

CV error (K=2): 0.48564

CV error (K=3): 0.36538

CV error (K=4): 0.28738

CV error (K=5): 0.22379

CV error (K=6): 0.21977

CV error (K=7): 0.19834

CV error (K=8): 0.22517

CV error (K=9): 0.21435

CV error (K=10): 0.26099

CV error (K=11): 0.27969
